## Supplementary information for "Energetic shifts reflect survival likelihood in *Anopheles gambiae*"

**<u>This document includes:</u>**

**Supplementary Table S1.** Effect of infection and blood meal on individuals that died naturally.
**Supplementary Table S2.** Spore load and resource dynamics in alive and dead mosquitoes across age classes.
**Supplementary Table S3.** Resource ratios dynamics in alive and dead mosquitoes across age classes.

**Supplementary Figure S1.** Average spore load in alive and dead mosquitoes.
**Supplementary Figure S2.** Resource degradation after different times post-death.
**Supplementary Figure S3.** Resource dynamics in alive and dead mosquitoes across age classes for infected and uninfected individuals.

**Supplementary Table S1. Effect of infection and blood meal on individuals that died naturally.** A linear model with longevity or spore load was used. As explanatory variables, we considered infection and blood meal in relation to the response variable longevity (Model 1a). For spore load (Model 1b), we considered age at death and blood meal (or not) as explanatory variables. In both cases, an interaction between the two factors was considered. Bold numbers reflect statistical significance.

| **Tested effect** | | | |
|---|---|---|---|
| **Model 1a** – Longevity | ***df*** | $\chi^2$ | ***p*** |
| Infection | 1 | 0.13 | 0.716 |
| Blood meal | 1 | 1.87 | 0.171 |
| Infection : Blood meal | 1 | 14.13 | **<0.001** |
| **Model 1b** – Spore load | ***df*** | $\chi^2$ | ***p*** |
| Age at death | 1 | 12.97 | **<0.001** |
| Blood meal | 1 | 3.91 | **0.048** |
| Age at death : Blood meal | 1 | 5.13 | **0.023** |

**Supplementary Table S2. Spore load and resource dynamics in alive and dead mosquitoes across age classes.** Summary of statistical tests for resource analyses. Protein, carbohydrate, and lipid contents were modelled with GLMs using appropriate error structures with blood meal, death status, infection, age at collection, and their interactions as explanatory variables.

| **Tested effect** | | | |
|---|---|---|---|
| **Model 2a** – Spore load | ***df*** | $\chi^2$ | ***p*** |
| Death status | 1 | 8.32 | **0.004** |
| Blood meal | 1 | 1.25 | 0.263 |
| Age class | 5 | 27.59 | **<0.001** |
| Death status : Blood meal | 1 | 1.87 | 0.172 |
| Death status : Age class | 5 | 3.64 | 0.602 |
| Blood meal : Age class | 5 | 14.18 | **0.014** |
| Death status : Blood meal : Age class | 5 | 4.47 | 0.483 |
| **Model 2b** – Proteins | ***df*** | $\chi^2$ | ***p*** |
| Death status | 1 | 15.15 | **<0.001** |
| Infection | 1 | 0.47 | 0.492 |
| Blood meal | 1 | 0.06 | 0.799 |
| Age class | 5 | 17.31 | **0.004** |
| Death status : Infection | 1 | 0.53 | 0.466 |
| Death status : Blood meal | 1 | 1.84 | 0.175 |
| Infection : Blood meal | 1 | 0.51 | 0.476 |
| Death status : Age class | 5 | 26.32 | **<0.001** |
| Infection : Age class | 5 | 2.32 | 0.803 |
| Blood meal : Age class | 5 | 5.54 | 0.354 |
| Death status : Infection : Blood meal | 1 | 1.00 | 0.318 |
| Death status : Infection : Age class | 5 | 2.03 | 0.844 |
| Death status : Blood meal : Age class | 5 | 7.04 | 0.217 |
| Infection : Blood meal : Age class | 5 | 1.89 | 0.864 |
| Death status : Infection : Blood meal : Age class | 5 | 2.93 | 0.710 |
| **Model 2c** – Carbohydrates | ***df*** | $\chi^2$ | ***p*** |
| Death status | 1 | 7.02 | **0.008** |
| Infection | 1 | 0.01 | 0.910 |
| Blood meal | 1 | 0.05 | 0.820 |

| | | | |
|---|---|---|---|
| Age class | 5 | 50.57 | **<0.001** |
| Death status : Infection | 1 | 0.89 | 0.344 |
| Death status : Blood meal | 1 | 0.12 | 0.728 |
| Infection : Blood meal | 1 | 0.01 | 0.915 |
| Death status : Age class | 5 | 34.21 | **<0.001** |
| Infection : Age class | 5 | 16.54 | **0.005** |
| Blood meal : Age class | 5 | 9.06 | 0.106 |
| Death status : Infection : Blood meal | 1 | 3.44 | 0.063 |
| Death status : Infection : Age class | 5 | 11.86 | **0.037** |
| Death status : Blood meal : Age class | 5 | 7.45 | 0.189 |
| Infection : Blood meal : Age class | 5 | 21.04 | **<0.001** |
| Death status : Infection : Blood meal : Age class | 5 | 22.42 | **<0.001** |
| **Model 2d** – Lipids | ***df*** | $\chi^2$ | ***p*** |
| Death status | 1 | 0.52 | 0.472 |
| Infection | 1 | 0.73 | 0.391 |
| Blood meal | 1 | 0.01 | 0.907 |
| Age class | 5 | 52.16 | **<0.001** |
| Death status : Infection | 1 | 5.17 | **0.023** |
| Death status : Blood meal | 1 | 2.23 | 0.135 |
| Infection : Blood meal | 1 | 0.25 | 0.619 |
| Death status : Age class | 5 | 24.18 | **<0.001** |
| Infection : Age class | 5 | 3.69 | 0.594 |
| Blood meal : Age class | 5 | 14.36 | **0.013** |
| Death status : Infection : Blood meal | 1 | 3.93 | **0.047** |
| Death status : Infection : Age class | 5 | 6.41 | 0.268 |
| Death status : Blood meal : Age class | 5 | 17.00 | **0.004** |
| Infection : Blood meal : Age class | 5 | 13.08 | **0.022** |
| Death status : Infection : Blood meal : Age class | 5 | 14.95 | **0.010** |

**Supplementary Table S3. Resource ratios dynamics in alive and dead mosquitoes across age classes**. Statistical analyses of resource ratios (P:C, P:L, C:L) in mosquitoes. Ratios were calculated after adding 1 to each value and analyzed with GLMs using a Gamma distribution of errors. Predictors included blood meal, death status, infection with Vavraia culicis, age at collection, and their interactions.

| **Tested effect** | | | |
|---|---|---|---|
| **Model 3a** – P:C | ***df*** | $\chi^2$ | ***p*** |
| Death status | 1 | 96.36 | **<0.001** |
| Infection | 1 | 0.10 | 0.751 |
| Blood meal | 1 | 0.21 | 0.644 |
| Age class | 5 | 199.24 | **<0.001** |
| Death status : Infection | 1 | 1.47 | 0.224 |
| Death status : Blood meal | 1 | 0.61 | 0.433 |
| Infection : Blood meal | 1 | 0.02 | 0.887 |
| Death status : Age class | 5 | 123.75 | **<0.001** |
| Infection : Age class | 5 | 81.18 | **<0.001** |
| Blood meal : Age class | 5 | 61.65 | **<0.001** |
| Death status : Infection : Blood meal | 1 | 0.95 | 0.330 |
| Death status : Infection : Age class | 5 | 7.85 | 0.165 |
| Death status : Blood meal : Age class | 5 | 6.21 | 0.286 |
| Infection : Blood meal : Age class | 5 | 128.88 | **<0.001** |
| Death status : Infection : Blood meal : Age class | 5 | 31.31 | **<0.001** |
| **Model 3b** – P:L | ***df*** | $\chi^2$ | ***p*** |
| Death status | 1 | 40.96 | **<0.001** |
| Infection | 1 | 0.04 | 0.838 |
| Blood meal | 1 | 0.01 | 0.909 |
| Age class | 5 | 191.11 | **<0.001** |
| Death status : Infection | 1 | 0.36 | 0.548 |
| Death status : Blood meal | 1 | 0.10 | 0.748 |
| Infection : Blood meal | 1 | 0.23 | 0.629 |
| Death status : Age class | 5 | 104.19 | **<0.001** |
| Infection : Age class | 5 | 3.75 | 0.586 |
| Blood meal : Age class | 5 | 84.52 | **<0.001** |
| Death status : Infection : Blood meal | 1 | 0.01 | 0.920 |

| | | | |
|---|---|---|---|
| Death status : Infection : Age class | 5 | 0.69 | 0.984 |
| Death status : Blood meal : Age class | 5 | 79.01 | **<0.001** |
| Infection : Blood meal : Age class | 5 | 7.82 | 0.166 |
| Death status : Infection : Blood meal : Age class | 5 | 5.42 | 0.366 |
| **Model 3c** – C:L | ***df*** | $\chi^2$ | ***p*** |
| Death status | 1 | 10.97 | **<0.001** |
| Infection | 1 | 0.02 | 0.890 |
| Blood meal | 1 | 0.12 | 0.726 |
| Age class | 5 | 153.87 | **<0.001** |
| Death status : Infection | 1 | 0.50 | 0.478 |
| Death status : Blood meal | 1 | 0.00 | 0.968 |
| Infection : Blood meal | 1 | 0.62 | 0.430 |
| Death status : Age class | 5 | 58.72 | **<0.001** |
| Infection : Age class | 5 | 13.46 | **0.019** |
| Blood meal : Age class | 5 | 24.36 | **<0.001** |
| Death status : Infection : Blood meal | 1 | 0.83 | 0.361 |
| Death status : Infection : Age class | 5 | 11.35 | **0.045** |
| Death status : Blood meal : Age class | 5 | 2.33 | 0.801 |
| Infection : Blood meal : Age class | 5 | 14.38 | **0.013** |
| Death status : Infection : Blood meal : Age class | 5 | 22.13 | **<0.001** |

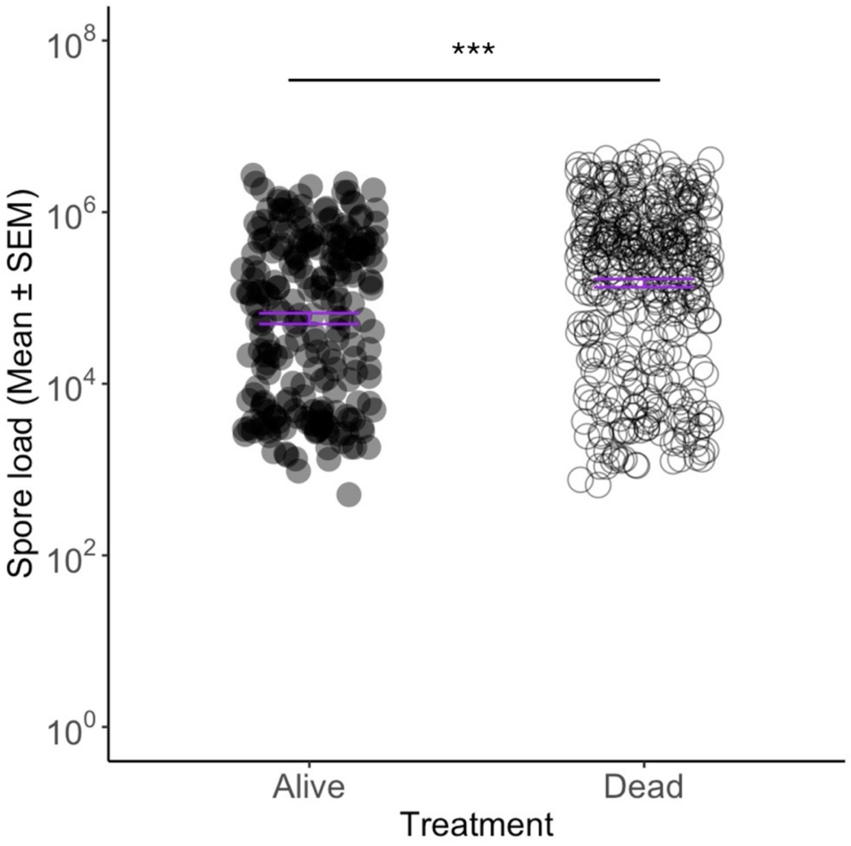


**Supplementary Figure S1. Average spore load in alive and dead mosquitoes.** Spore load was higher in dead than alive mosquitoes (dead: $5.0 \times 10^5$; alive: $2.1 \times 10^5$; $\chi^2 = 8.32$, df = 1, p = 0.004), independently of their age (interaction death status * age: $\chi^2 = 3.64$, df = 5, p = 0.602). The error bars represent the standard error of the mean (SEM), and an asterisk indicates statistically significant differences between alive and dead individuals. The sample sizes (*n*) were 419 dead and 231 alive**.**

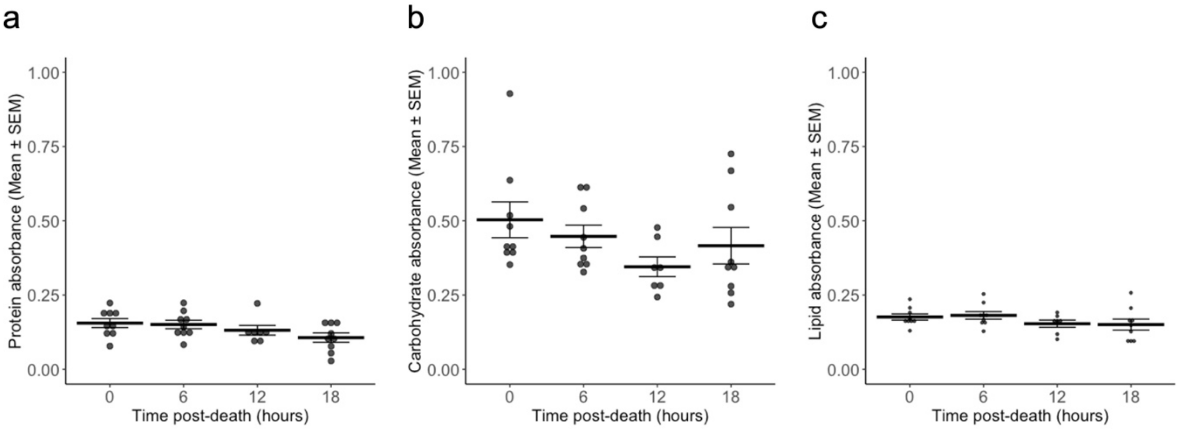


**Supplementary Figure S2. Resource degradation at different times after death.** A preliminary experiment was carried out to estimate if there is resource degradation at different time points post mosquito death: 0, 6, 12, and 18 hours post-death. For this, eight three-day-old mosquitoes (four exposed to *V. culicis* and four unexposed) per time-point were snap-frozen at -80 ºC for 24 hours and then left at room temperature for the duration of their time treatment. At the respective time-point, each individual was frozen at -80 ºC and their resources were quantified the next morning (see the Materials and Methods). The average protein (a), carbohydrate (b) and lipid (c) content for each of the time post-death treatments is plotted above using absorbance as a proxy. The error bars show the standard error of the mean (SEM). A linear model was run for each resource category, with resource content as the response variable and time as an explanatory variable. There was no significant effect of time on the average content for any of the resource classes (Proteins: $F_{3,30} = 2.197$, $p = 0.109$; Carbohydrates: $F_{3,30} = 1.508$, $p = 0.233$; Lipids: $F_{3,30} = 1.260$, $p = 0.306$).

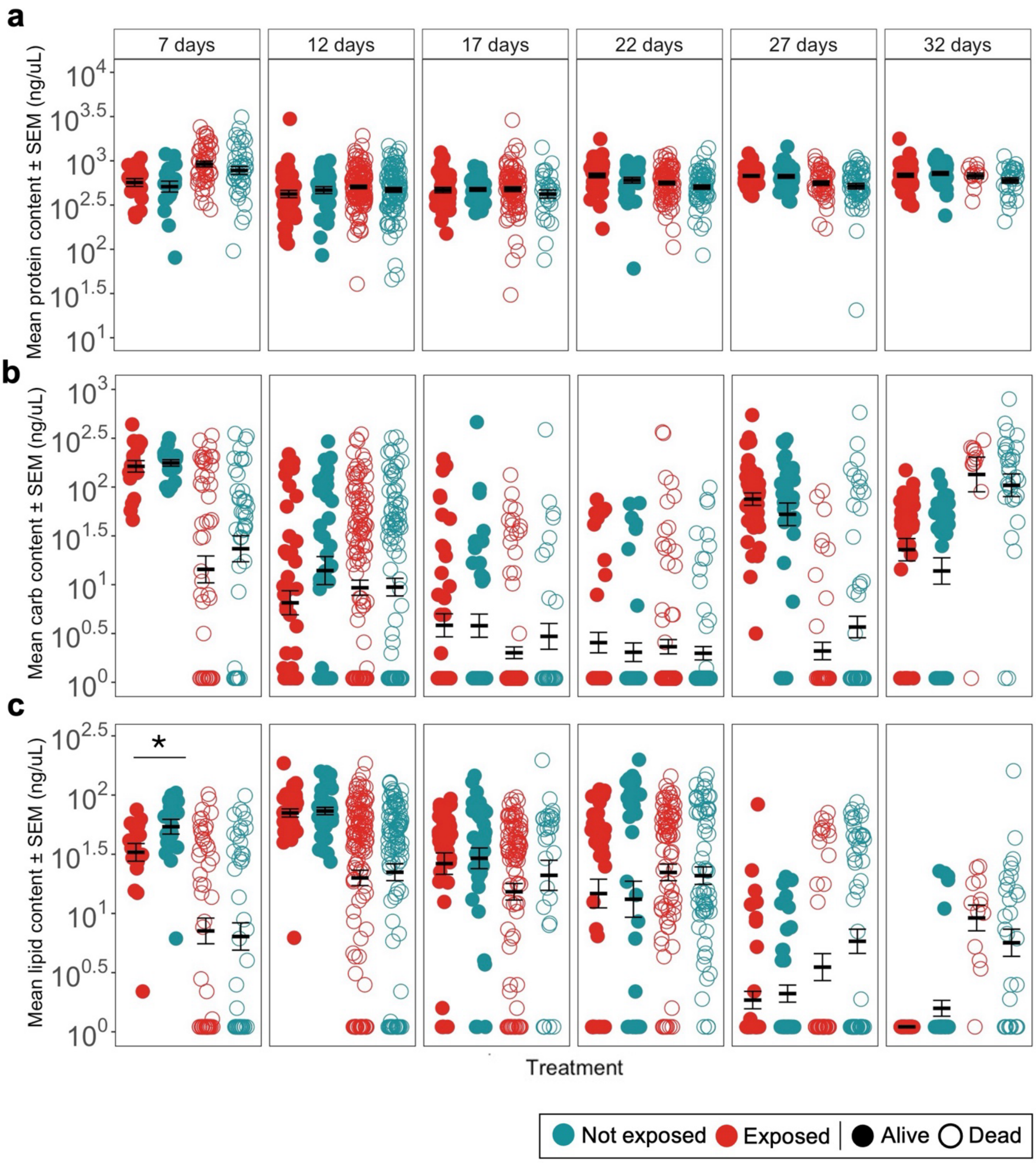


**Supplementary Figure S3. Resource dynamics in alive and dead mosquitoes across age classes for infected and uninfected individuals.** **(a)** Protein, **(b)** carbohydrate and **(c)** lipid content throughout their lifetime for infected and uninfected individuals that were collected alive or dead. The error bars show the SEM, and an asterisk indicates statistically significant differences from multiple comparisons between uninfected and infected individuals. The sample sizes (*n*) were 19, 20, 48 and 42 for day 7; 42, 36, 123 and 101 for day 12; 41, 39, 92 and 31 for day 17; 41, 39, 80 and 63 for day 22; 41, 37, 43 and 60 for day 27; and lastly 41, 38, 13 and 30 for day 32 in the treatment sequence of the caption for **(a)**. The sample sizes were 20, 20, 51 and 44 for day 7; 42, 38, 126 and 101 for day 12; 42, 39, 94 and 31 for day 17; 41, 39, 82 and 64 for day 22; 41, 39, 43 and 60 for day 27; and lastly 41, 39, 13 and 30 for day 32 in the treatment sequence of the caption for **(bc)**.
